# Phase separation induced by cohesin SMC protein complexes

**DOI:** 10.1101/2020.06.13.149716

**Authors:** Je-Kyung Ryu, Celine Bouchoux, Hon Wing Liu, Eugene Kim, Masashi Minamino, Ralph de Groot, Allard J. Katan, Andrea Bonato, Davide Marenduzzo, Davide Michieletto, Frank Uhlmann, Cees Dekker

## Abstract

Cohesin is a key protein complex that organizes the spatial structure of chromosomes during interphase. Here, we show that yeast cohesin shows pronounced clustering on DNA in an ATP-independent manner, exhibiting all the hallmarks of phase separation. *In vitro* visualization of cohesin on DNA shows DNA-cohesin clusters that exhibit liquid-like behavior. This includes mutual fusion and reversible dissociation upon depleting the cohesin concentration, increasing the ionic strength, or adding 1,6-hexanediol, conditions that disrupt weak interactions. We discuss how bridging-induced phase separation can explain the DNA-cohesin clustering through DNA-cohesin-DNA bridges. We confirm that, *in vivo,* a fraction of cohesin associates with chromatin in yeast cells in a manner consistent with phase separation. Our findings establish that SMC proteins can exhibit phase separation, which has potential to clarify previously unexplained aspects of *in vivo* SMC behavior and constitute an additional principle by which SMC complexes impact genome organization.

**One sentence summary:** Yeast cohesin complex is observed to phase separate with DNA into liquid droplets, which it accomplishes by ATP-independent DNA bridging.

---

Members of the structural maintenance of chromosome (SMC) protein family such as condensin, cohesin, and the Smc5/6 complex are key proteins for the spatial and temporal organization of chromosomes (*1–4*). Recent *in vitro* experiments visualized real-time DNA loop extrusion mediated by condensin and cohesin (*5–7*). While loop extrusion by SMC proteins constitutes a fundamental building block in the organization of chromosomes, other factors may also contribute. In the last decade, it has become abundantly clear that phase separation plays a role in many processes in biological cells (*8*), including chromosome organization (*9,10*). Thus far, SMC proteins have not been implied in phase separation. While ATP-independent clustering of DNA and SMC proteins has been reported (*11–14*), such observations were attributed to imperfect protein purification or non-physiological buffer conditions. For example, Davidson et al. (*7*) reported *in vitro* DNA loop extrusion by the human cohesin complex when the complex concentration was limited to very low values (<0.8 nM, i.e., much lower that physiological concentrations of ~333 nM (*15, 16*), and mentioned that the cohesin complexes were prone to aggregation at higher concentrations. Such findings raise the question whether such aggregate formation may be intrinsic and may have a physiological meaning.

Here we report that interactions between the yeast cohesin SMC complex and DNA lead to pronounced phase separation. This clustering behavior is ATP independent but depends on DNA length. We find that single cohesin complexes are able to bridge distant points along DNA that act as nucleation points for recruiting further cohesin complexes – a behavior indicating bridging-induced phase separation (BIPS) (*17, 18*), also known as polymer-polymer phase separation (PPPS) (*19*), a type of phase condensation that was studied theoretically but lacked experimental verification so far.

First, we visualized cluster formation by *Saccharomyces cerevisiae* cohesin complex on DNA *in vitro* in real time (Fig. 1A and Movie S1). We immobilized SYTOX-Orange (SxO) labelled double-tethered λDNA (48.5 kbp) on a polyethylene glycol (PEG)-coated surface, and applied 10 nM cohesin holocomplexes (i.e. the cohesin tetramer Smc1-Smc3-Scc1-Scc3 plus the cohesin loader Scc2-Scc4). Note that these cohesin complexes are proficient in cohesin loader-stimulated ATP hydrolysis and topological loading onto DNA (*7, 20*) (fig. S1). We tested these yeast cohesin complexes extensively for their putative loop-extrusion activity, but we failed to observe any DNA-loop-extrusion activity over a very wide range of parameters and combinations of cohesin subunits. Instead, we observed the spontaneous accumulation of DNA spots along DNA molecules (Fig. 1A and Movie S1). Application of an in-plane side flow (*5*) showed that these were stably condensed clusters (Fig. 1B and Movie S2) and not DNA loops. Importantly, we observed clusters formed by cohesin complexes both in the absence and presence of ATP, showing that this behaviour is ATP independent. To quantify the kinetics of cluster formation, we measured the fluorescence intensity of the cluster region (Materials and methods) (*5*). Upon flushing in cohesin, the intensity at the cluster spot increased approximately linearly over time (Fig. 1C). After a compaction time of about 30 s, a plateau was reached, where the cluster comprised of a sizeable amount of DNA (20 ± 8 kbp; for more examples see fig. S2). As Figure 1D shows, cluster formation did not occur for cohesin tetramers (i.e., without the loader) within 400 s at 10 nM, although it was observed for higher concentrations (data not shown). Clustering proceeded slightly slower (~27 s) at room temperature than at 32 °C (~14 s). We conclude that the observed cluster formation is induced by the ATP-independent interactions between the cohesin complex and DNA.

**Figure 1.**
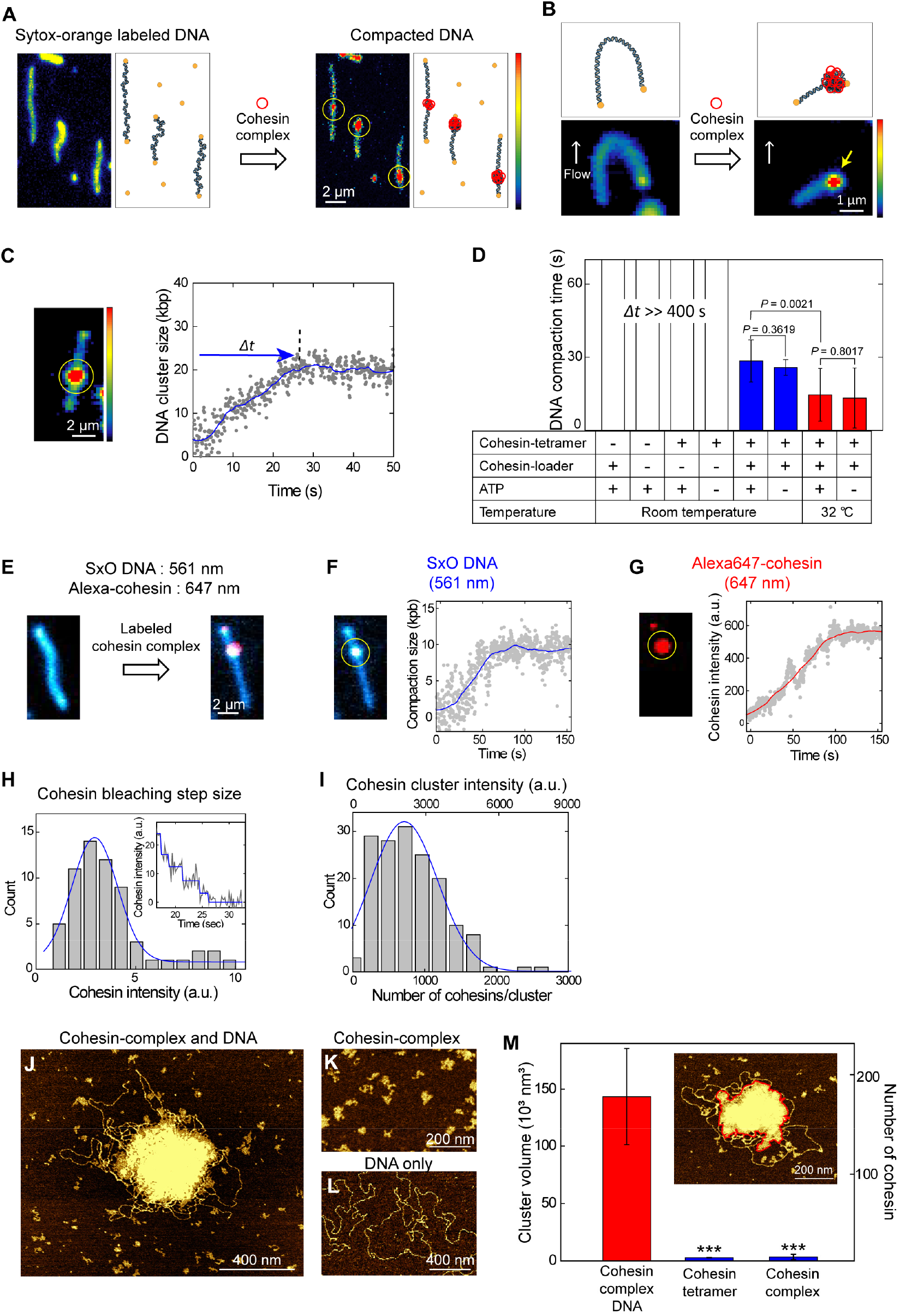
ATP-independent DNA compaction mediated by cohesin complex. (A) Snapshots before and after cohesin-induced compaction of a doubly tethered DNA molecule. In schematics on the right, blue represents DNA, yellow biotin-streptavidin, and red the cohesin complex. (B) Snapshots and schematics of side-flow experiment before and after addition of cohesin complexes. Flow is applied in both cases (white arrows). The yellow arrow indicates a DNA region that is tightly clustered (i.e. not a DNA loop). See also Movie S2. (C) DNA cluster size as a function of time (right) calculated from the integrated fluorescence intensities in the cluster and the full 48.5-kbp λDNA in the image (left). Yellow circle indicates the compaction spot. See also movie S3. (D) Comparison of the DNA compaction time at various conditions (*N* = 33, 25, 10, 13, 10, 13, 11, 9, respectively). Two paired Student *t*-test was used. (E) Cluster formation with labeled cohesin complex showing the co-localization of compacted DNA (blue) and cohesin (red). (F) Representative measured trace of DNA compaction. (G) Simultaneously measured trace for the cohesin binding. (H) Histogram of bleaching-step intensities of single Alexa647-cohesin molecules (*N* = 64). Inset shows a representative bleaching trace of a small cohesin-complex cluster with a low number of cohesin complexes along the DNA. A Gaussian fit (blue) yielded 3.0 ± 1.2 a.u. (mean ± SD). (I) Number of cohesin complexes in a cluster. A Gaussian fit (blue) yielded 740 ± 500 (mean ± SD). (J-L) AFM images of cohesin-complex/DNA, cohesin-complex only, and DNA only, respectively. (M) Volumes of the DNA/cohesin clusters for different conditions (median ± SEM; *N* = 21, 11, 57). Red lines in inset illustrates a cluster with its boundary (red). *** indicates *P* < 0.001 assessed by the two-paired Student *t*-test.

The clusters contained a large number of cohesin complexes, which we quantified by coimaging Alexa647-labelled cohesin complexes and SxO-labeled DNA. Cohesin complexes were observed to co-localize with the DNA-clusters (Fig. 1E and Movie S3) while hardly any cohesin was observed at other locations on the DNA. We consistently observed a simultaneous increase of both DNA (Fig. 1F) and cohesin intensity (Fig. 1G) in the clusters. To count the number of cohesin complexes on the DNA, we compared the cohesin intensities of each cluster (Fig. 1I, top axis) with the intensity of single cohesin complexes as deduced from bleaching steps in traces (Fig. 1H) yielding an estimate of 720 ± 470 (mean ± SD) cohesin complexes within a cluster (Fig. 1l, bottom axis). Next, we visualized the clusters at higher resolution using AFM imaging of mixtures of 4 ng/μL λDNA and 10 nM cohesin holocomplexes. Again, large cohesin-complex/DNA clusters were observed when both cohesin and DNA were present (Fig. 1, J and M), while no cluster formation was observed for cohesin-complex only, or DNA only (Fig. 1, K and L). The AFM images showed clusters with a dense protein-rich center, and an outer region made of loosely compacted loops. These cohesin-complex/DNA clusters contained many cohesin holocomplexes in a broad distribution with a median value of about 170, a number that was estimated by dividing the average cluster volume by the volume of an individual cohesin complex (Fig. 1M and fig. S3). Both the fluorescence and AFM data thus indicate a very large number of cohesin complexes per cluster (where the lower estimate from AFM likely originates from the lower protein/DNA ratio). These results show that cluster formation is not due to DNA-independent oligomerization of cohesin complexes but instead relies on interactions between cohesin complexes and DNA.

Strikingly, the cohesin-DNA clusters displayed the behavior of liquid droplets. The clusters exhibited an average size of 1.14 ± 0.18 μm that clearly exceeded the size of diffraction-limited spots measured for 20 nm quantum dots (0.57 ± 0.11 μm) (Fig.2, A and B). The droplets were spherical in shape (Fig. 2C). When multiple clusters formed on a single DNA molecule (fig. S4), we often observed that two spherical neighboring clusters merged over time, where subsequently the shape of the resulting cluster again became spherical (Fig. 2, D and E, and Movie S4). These features can be viewed as a defining behavior of liquid droplets and direct evidence of phase condensation.

**Figure 2.**
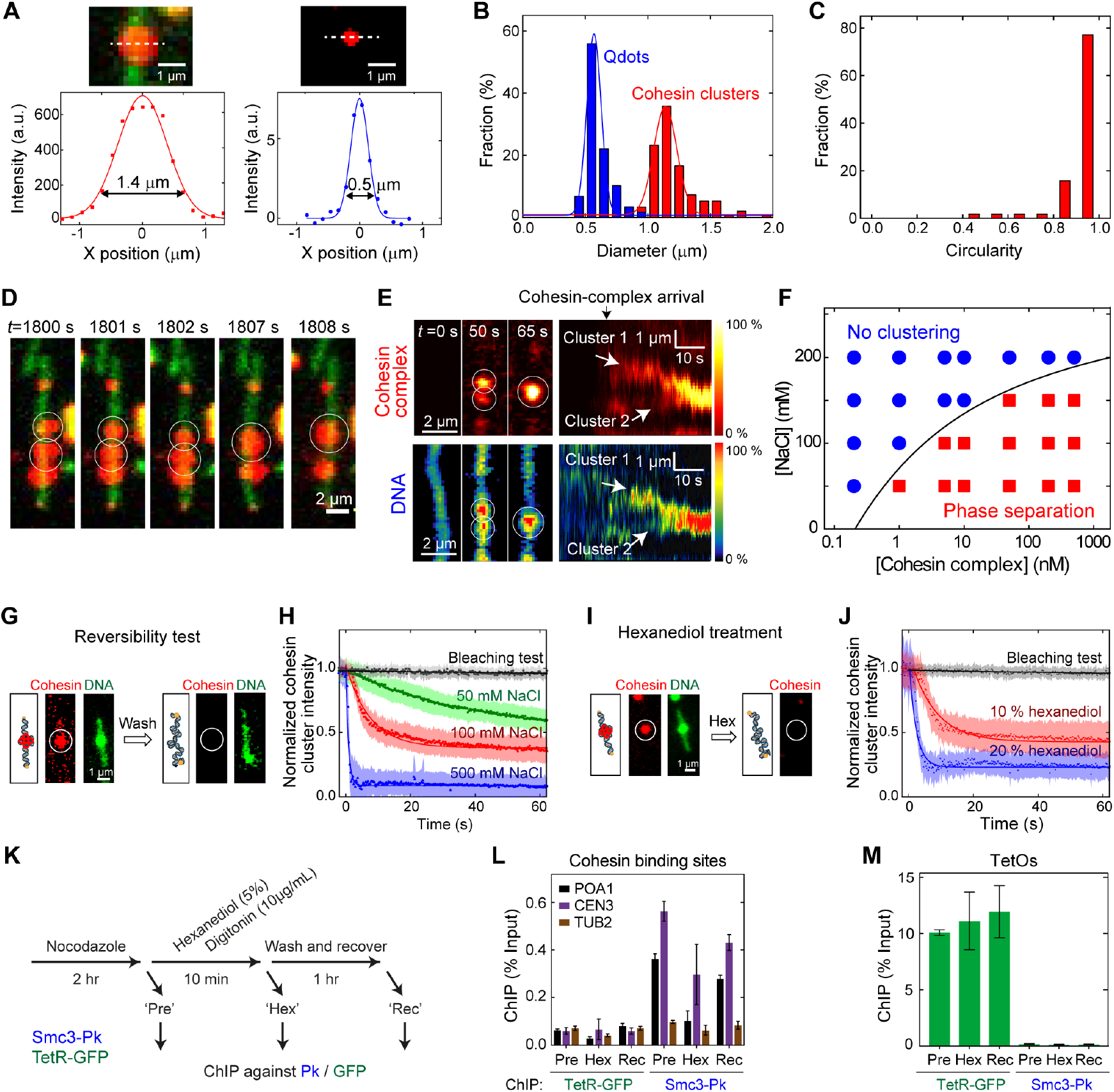
Cohesin complex forms liquid droplets along DNA *in vitro* and *in vivo.* **(A)** Images of a cohesin-complex/DNA droplet (top left) and 20 nm QD (top right). White lines indicate where cross-sectional intensity profiles were acquired, yielding the bottom panels with Gaussian fits. Cluster size was measured as the width at 20 % of the maximum fluorescence intensity of the profile. **(B)** Diameter distributions of cohesin-complex droplets (mean ± SD = 1.14 ± 0.18 μm, *N* = 151) and QD (mean ± SD = 0.57 ± 0.11 μm, *N* = 59). **(C)** Circularity distributions of cohesin droplets (*N* = 57). The circularity was measured as *4πA/P*^2^, where *A* and *P* are the area and perimeter of the droplet, and it shows how closely the shape resembles a perfect circle with a circularity of 1 (*9*). **(D)** Cohesin complex (red) forms liquidphase droplets along a DNA (green). Over time, two droplets are seen to fuse into one spherical droplet (Movie S4). **(E)** Merging of two clusters, as monitored in the cohesin (top) and DNA (bottom) channels. Left show three snapshots; rights shows fluorescence intensity kymographs (*N* = 18). **(F)** Phase diagram of cluster formation induced by cohesin complex and λDNA at various salt (NaCl) and cohesin-complex concentrations. Red squares indicate that DNA compaction occurred, while blue circles indicate that this did not occur. Solid line is a guide to the eye. **(G)** Reversibility test that shows the dissolving of a cluster by washing in a high salt buffer (500 mM). **(H)** Cohesin cluster intensities versus time upon washing with buffers with different salt concentrations. Light-colored areas denote the SD of the data points. Solid lines are exponential fits yielding dissociation times of 476 ± 25 s, 41.9 ± 0.7 s, 7.5 ± 0.2, and 0.79 ± 0.03 s (errors are SD) for bleaching, 50, 100, and 500 mM NaCl, respectively (*N* = 60, 113, 93, 150). **(I)** Dissolving of a cohesin cluster by washing with hexanediol. **(J)** Cohesin cluster intensity versus time upon washing with hexanediol. Lightcolored areas denote the SD of the data points. Dissociation times are 476 ± 25 s, 7.9 ± 0.1 s and 2.2 ± 0.1 s for bleaching, 10 %, and 20 % 1,6-hexanediol buffer, respectively (*N* = 60, 80, 37). **(K)** Scheme for *in vivo* hexanediol treatment of yeast cells. **(L)** Chip-qPCR results for cohesin (Smc3-Pk) and binding to a centromere (*CEN3*), chromosome arm (*POA1*), and a negative-control binding site (*TUB2*), before (Pre) and after hexanediol treatment (Hex), and after washing out hexanediol (Rec) (mean ± SEM, *N* = 3). TetR-GFP binding was analysed as a control. (M) Same for binding to tetO sites (mean ± SEM, *N* = 3).

Another characteristic feature of phase condensates is the reversibility of their formation. Cohesin-DNA clusters could be dissolved upon depleting the cohesin-complex concentration in the buffer, or by increasing its salt concentration – thereby demonstrating the reversibility of the phase condensation. A phase diagram of cluster formation shows that clustering is favored by low salt and high cohesin concentrations (Fig. 2F and fig. S5) – a feature observed for many phase-separating proteins (*9*). Importantly, cluster formation was observed at physiologically relevant concentrations of salt (~150 mM NaCl) and cohesin (> 1 μM in yeast (*15*); 333 nM in human (*16*)), suggesting that cluster formation may as well occur *in vivo.* When the cohesin complex was depleted by washing the channel with buffer, we observed the dissociation of the cohesin-DNA clusters as a decrease in the intensities of the cohesin-complex droplets on DNA (Fig. 2, G and H). This implies that the clusters are dynamic, i.e., cohesins in clusters exchange with the pool in the bulk solution. The dissociation of cohesin-complex clusters was fastest at elevated salt concentrations, indicating that electrostatic interactions underlie droplet formation (Fig. 2H).

Next, we used 1,6-hexanediol, an aliphatic alcohol that interferes with weak protein-protein (*21*) and protein-nucleic acid interactions (*22*) and is often used to differentiate liquid-phase and solid-like biological condensates as it dissolves liquid droplets but not gel-phase assemblies (*9, 21*). *In vitro*, we observed that 1,6-hexanediol disrupted cohesin-DNA droplets (Fig. 2, I and J). Subsequent addition of cohesin complexes in a buffer without hexanediol led to recovery of DNA-cohesin droplets (fig. S6). Furthermore, we transiently applied 1,6-hexanediol to live yeast cells (Fig. 2K). Cohesin levels on chromosomes were monitored by chromatin immunoprecipitation (ChIP) followed by quantitative real-time PCR (qPCR). Ten minutes after administering 1,6-hexanediol, the amount of cohesin-associated DNA had noticeably decreased, which recovered again following 1,6-hexanediol washout (Fig. 2L). This suggests that a portion of cellular cohesin reversibly associates with chromosomes through phase separation, in addition to cohesin that topologically embraces DNA (*23*). To confirm that 1,6-hexanediol had not nonspecifically disrupted DNA-protein interactions, we monitored the association of a tetracyclin repressor-GFP (tetR-GFP) fusion protein with tetracyclin operators (tetOs) contained in the same strain. Its chromatin binding remained unaltered upon 1,6-hexanediol treatment (Fig. 2M), indicating that DNA binding of cohesin *in vivo* is uniquely susceptible to an agent that disrupts phase condensates.

Unexpectedly, the cohesin-DNA cluster formation critically depended on DNA length (Fig. 3 A-F). For short DNA lengths *l*, the cluster size, characterized by its radius of gyration *R_G_*, remained insensitive to *l* (Fig. 3E, blue line), whereas beyond a critical value of *l_c_* ≈ 3 kbp, we observed that the cluster size strongly increased with DNA length, scaling as a power law *R_G_*~ *l^α^*, with *α* = 0.45 ± 0.01 (Fig. 3E, red line; error is SD). The length-independent cluster size at short DNA length can simply be attributed to the size of single cohesin-complex binding to DNA (Fig. 3, A and F). Importantly, the absence of any clustering for short DNA indicates that the cohesin binding to DNA does not trigger any significant cohesin-cohesin interaction, e.g., through some conformational change. The critical value *l_c_* ≈ 3 kbp that marks the onset of significant length-dependent clustering is in remarkably agreement to the length of DNA for which thermal fluctuations can induce spontaneous looping, i.e., *l_c_ = 2π^2^l_P_* ≅ 987 nm ≅ 2903 bp for a persistence length *l_P_ =* 50 nm (see SI). Such stochastic thermal loops can be stabilized by a cohesin complex that subsequently acts as a nucleation point for a cluster via BIPS (*17, 18*), whereupon further clustering is entropically and energetically favored over a dispersed cohesin distribution.

**Figure 3.**
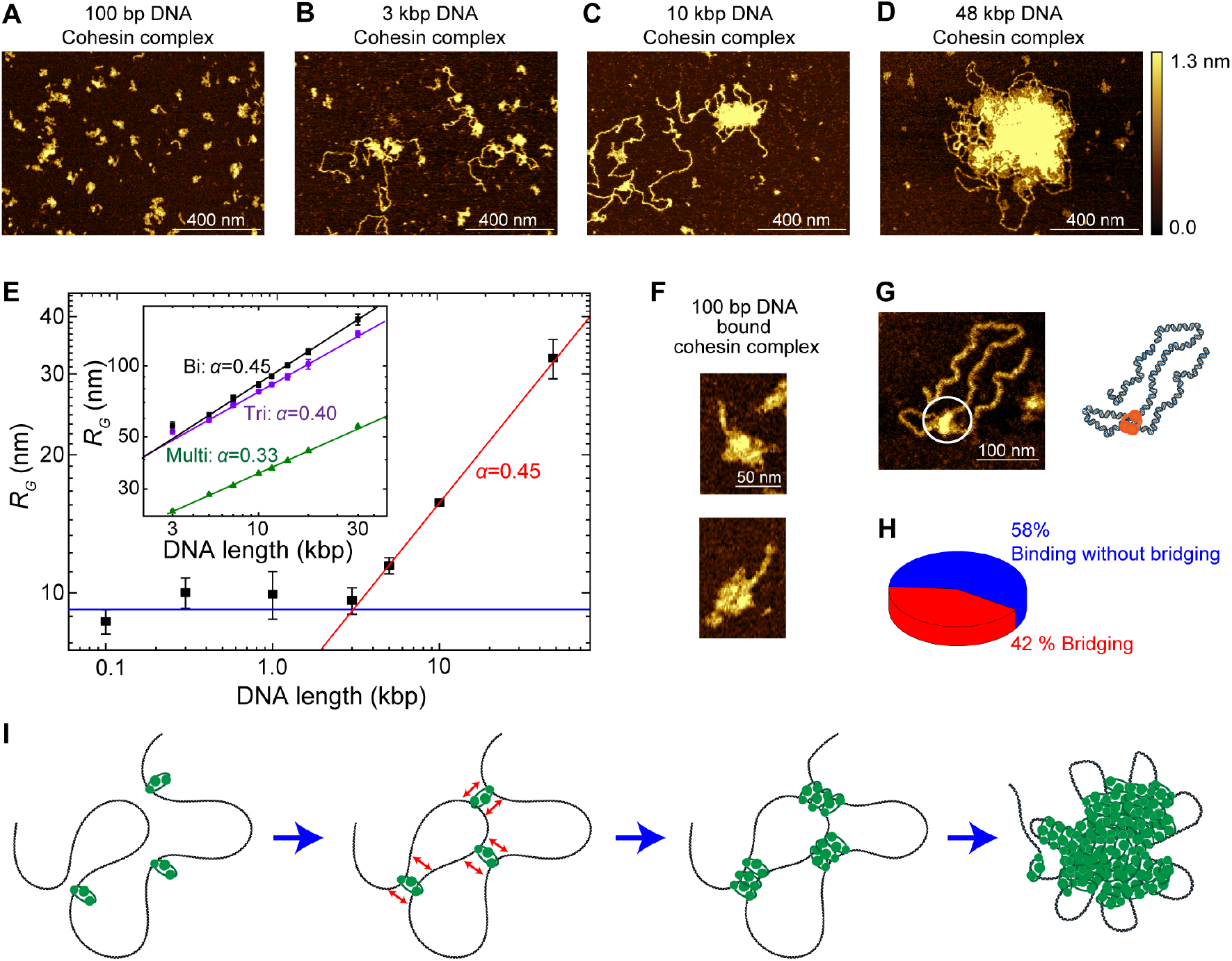
AFM imaging of DNA-mediated cohesin clusters, and BIPS model. **(A-D)** Representative AFM images of cohesin complex as it binds to different length DNA of 100 bp, 3 kbp, 5 kbp, and 48.5 kbp, respectively. **(E)** Radius of gyration of cohesin-DNA clusters versus DNA length. Note the log-log scale. Error bars are SEM. At low DNA length *l*, the cluster size is constant (green line), while beyond a critical length *l_c_* ≈ 3 kbp, the data exhibit a power law, *R_G_* ~ *l^α^* (blue line) with *α* = 0.45 ± 0.01 (SD) from a fit. Inset: Data from MD simulations for cohesin with bi-(black, *α* = 0.45 ± 0.01), tri-(violet, *α* = 0.40 ± 0.01), and multi-connectivity (green, *α* = 0.33 ± 0.01). **(F)** Examples of single cohesin complex bound to 100 bp DNA. **(G)** Representative image of DNA bridging by a single cohesin complex. **(H)** Probability that a single cohesin complex that bound DNA did or did not bridge to another segment along the DNA (*N* = 81). **(I)** Working BIPS model for cohesin-mediated phase separation. Bare DNA is bridged by a single-cohesin complex, increasing the local concentration of DNA, which subsequently induces binding of more cohesin complexes to this region, leading to the formation of a large DNA/cohesin-complex droplet.

The power-law scaling of cluster size with DNA length reveals underlying properties of the condensation. While a power-law scaling with *α* = 1/3 is generally associated with collapsed globular polymer conformations, we here find a higher exponent of *a =* 0.45, closer to that of an ideal polymer (*α* = 0.5) (*24*). We performed Molecular Dynamics (MD) simulations in which DNA-binding bridges are modelled as patchy particles with 2, 3, or many (~10) binding sites (see Fig. 3E, inset, fig. S7, and Movie S5). As expected, we found that the many-binding case led to the formation of a compacted globule (fig. S7C) with *α* = 0.33. By contrast, clusters formed by cohesin bridges with *n* = 2 or 3 binding sites induced a qualitatively different condensation with an exponent of 0.45 and 0.40, respectively (Fig. 3E, inset and fig. S7D), with the *n* = 2 result in excellent agreement with our AFM data. Interestingly, the MD simulations for lower *n* showed cohesin-rich droplets that were rather porous and penetrable to diffusing solutes of sufficiently small size. Notably, while cohesin models with many (~10) binding points induced, as expected (*18*), Hi-C checker-board patterns that are qualitatively similar to mammalian compartments (fig. S8) (*10*), bridges with only 2 or 3 binding sites induced only very weak long-range compartments in *in silico* HiC maps, in line with experimental data for budding yeast cohesin that show only weak compartmentalization (*25, 26*).

Our results suggest that a BIPS model (Fig. 3I) underlies DNA-mediated cohesin phase separation. In such a scenario, formation of droplets is initiated via cohesin complexes that bridge the DNA polymer (*19*), whereupon additional proteins bind near the first bridging sites, in turn yielding larger clusters (*17*). This process is driven by a positive feedback between bridging and local DNA concentration (*18*). Local bridging of distant segments along a DNA molecule is an essential element in BIPS. Using AFM imaging, we indeed observed that single cohesin complexes can bridge the DNA, implying (at least) two DNA binding sites within the complex, consistent with previous suggestions of multiple potential binding sites in budding yeast cohesin (*27*). Figure 3G is an example where a single cohesin complex is seen to bridge DNA (see fig. S9 for more examples), which was obtained by incubating 3 kbp DNA and cohesin complexes for a very short time. Here, over 40 % of single cohesin complexes that bound to DNA showed DNA bridge formation (Fig. 3H).

Our observations of cohesin phase separation may help to interpret previous unsolved questions about genome organization by SMC proteins. First, SMC-mediated phase separation explains why many previous *in vitro* studies observed higher-order assemblies with DNA and SMC proteins in the absence of ATP (*11–14*). Second, *in vivo,* yeast cohesin has been observed to exist in foci on spread chromosomes (*28*), consistent with phase separated clusters. Interallelic complementation between two mutant alleles in the same cohesin subunit has provided functional evidence for as yet unexplained cohesin-cohesin interactions (*29*). Phase separation could explain such interactions as well as *in vivo* chromatin recruitment of cohesin complexes that are unable to topologically embrace DNA on their own (*23*). Third, phase-separation induced by cohesin complexes may explain how DNA loops can be stabilized at CTCF bound sites in vertebrates. As CTCF is unstably bound to DNA (residence time 1-2 min) and cohesin is more stably bound (residence time 22 min) (*30*), cohesin phase separation at the stem of loops could stabilize cohesin-CTCF complexes. And finally, our simulations showed that cohesin phase separation only yielded weak long-range compartment patterns which notably is consistent with experiments that showed stronger A/B compartmentalization upon cohesin depletion (*10*).

Summing up, the demonstration that cohesin is a phase-separating enzyme reveals another basic principle for organizing genome architecture that potentially may be a generic feature of other SMC proteins as well.

## ACKNOWLEDGMENTS

We thank J. van der Torre for technical support, and thank R. Barth, J.-M. Choi, J. Eeftens, M. Ganji, J. Kerssemakers, S. Kim, S.H. Kim, B. Pradhan, L. Reese, S.H. Rah, E. van der Sluis, and H. C. Treurniet for discussions.

## Author contributions

J.-K. R., F. U. and C. D. designed the experiments, J.-K. R., E. K., and R. G. performed single-molecule visualization assay, J.-K. R. and A. J. K. performed AFM experiments, C. B., M. M, and F. U. purified cohesin proteins, H. W. L performed *in vivo* hexanediol experiments, D. Mi. designed the theory and computational model, A.B, D. Mi., and D. Ma. performed MD simulations, J.-K. R. and R. G. performed ATPase assays, J.-K. R., F. U., and C. D. wrote the manuscript, and all authors provided input to that.

## Data and materials availability

All data are available in the manuscript or the supplementary material, or upon request.

## Supplementary Materials

### Materials and Methods

#### Protein purification

Cohesin tetramer (Smc1, Smc3, Scc1, and Scc3) and cohesin loader (Scc2-Scc4) were purified by following protocols described in a previous paper (*20*). For the labelling of cohesin, integrative plasmids containing pGAL-SMC1-PK3 ADE2, pGAL-OptSCC1-HIS-3C-ProtA, pGAL-SMC3-SNAP TRP and pGAL-SCC3-My URA3 were transformed (strain Y5345) to obtain SNAP-fused Smc3 in C-termini. For this construct, we used the same purification protocol.

#### Fluorescent labelling of cohesin

We mixed 1.16 μM SNAP-tag cohesin tetramer with 40 μM SNAP-Surface Alexa Fluor 647 (NEB) in a 20 mM Tris-pH7.5, 150 mM NaCl, 0.05 %(w/v) Tween20, 3 %(w/v) Glycerol, 0.1 mg/mL BSA, 1 mM DTT, and incubate the mixtures for an overnight. Labelled protein was separated from free fluorophore using a Zeba Microspin 40 kDa (Thermo Fisher), we filtered free fluorophores in 20 mM Tris, 150 mM NaCl, 0.05 % Tween20, and 3 % Glycerol.

#### Preparation of biotin-labelled λDNA

Lambda DNA was labelled with biotin at both ends following protocols in a previous paper (*5*).

#### ATP hydrolysis assay

A high-throughput colorimetric ATPase assay (PiColorLock, Expedeon) was used to measure the ATPase activity of cohesin complex by following the manufacturer’s protocol. ATPase reactions were set up in total volumes of 20 μL containing with 40 mM Tris-pH 7.5, 25 mM NaCl, 2.5 mM MgCl_2_, 2.5 mM ATP., and 0.5 mM TCEP. Concentrations of 100 ng/μL λDNA (Promega) and 50 nM cohesin with or without 50 nM loader were used. Reactions were initiated by the addition of cohesin complex and incubated for 15 min. Temperature was controlled using a a PCR machine. ATP hydrolysis was halted by adding 5 μL PiColorLock^TM^ reagent into the reaction mixture which also initiated color development. After 2 min, a stabilizer was added and thoroughly mixed to stop the coloration. Using a nanodrop spectrometer, the absorbance was measured at 640 nm. The hydrolysis rate was calculated by dividing the total amount of phosphate that was produced by the total reaction time.

#### Single molecule fluorescence assay

Microfluidic flow chambers for fluorescence imaging were prepared by following an established protocol (*31*). Chamber dimensions were 3 mm × 15 mm × 100 μm. Piranha was used to clean a quartz slide to which 12 holes were drilled. The quartz slide and cover slips were PEGylated with a 1-to-100 ratio of biotin-PEG and PEG by dissolving the powders into sodium bicarbonate solution (0.1 M NaHCO_3_, pH 8.5). After washing and drying the PEGylated slides, the flow chambers were assembled using double-sticky tape that defined 6 chambers, and the edges of the chambers were sealed by epoxy glue. Polytetrafluoroethylene (PTFE) tubing was connected to the drilled holes on one side of each chamber and a reservoir was built using a pipette tip.

Buffer flow was controlled with an electric syringe pump. First, T50 buffer (20 mM Tris-pH7.5, 50 mM NaCl) was injected into the flow channel. In order to tether the DNA onto the PEG surface, 100 μg /mL streptavidin solution was flushed for 30 sec in order for streptavidin to bind to the biotin-PEG, followed by washing with T50 buffer. A solution of 48.5 kbp double-biotinylated λDNA (100 pg/μl) in T50 buffer was injected at an initial flow speed of 20 μL/min for 30 seconds. After that, a reduced flow speed of 3 μL/min was maintained for 20 minutes. Unless stated otherwise, single-molecule fluorescence cohesin studies were performed in a reaction buffer of 50 mM Tris-pH 7.5, 50 mM NaCl, 2.5 MgCl_2_, 0.5 TCEP mM, 0.5 mg/mL BSA, 10-50 nM SYTOX-Orange (SxO), and 2.5 mM ATP. Most experiments were done using 10 nM cohesin complexes (cohesin tetramer and cohesin loader). Lower concentration of SxO was used to minimize labelling artefacts. To observe the phase diagram of Fig. 2C, we used a variety of NaCl and cohesin-complex concentrations. For experiments with labelled cohesin, an imaging buffer was used of 100 mM Tris-pH7.5, 50 mM NaCl, 2.5 mM MgCl_2_, 50 nM SYTOX-Orange, and the oxygen scavenging system (2 mM Trolox, 1 % glucose, 300 μg/mL glucose oxidase, 30 μg/mL catalase).

For imaging of the SxO-stained DNA only, a 561 nm laser excitation was used. For dualcolor colocalization imaging of SxO-stained DNA and Alexa647-labelled cohesin, an alternating laser excitation mode (ALEX) was used with 561 nm and 642 nm laser excitation, respectively. We used a custom-modified inverted Nikon epifluorescence microscope equipped with a Nikon 100x / 1.49 Apo TIRF oil immersion objective. Image acquisition started immediately after injection of the reaction buffer. Highly inclined and laminated optical sheet (HILO) mode was used for imaging, and temperature was controlled (Okolab). Images where acquired by a CCD camera (Andor iXon Ultra 897) with a dual emission image splitter (OptoSplit) for dual-color experiments. Metamorph software was used to record the singlemolecule fluorescence images.

#### In vitro high-salt wash and hexanediol treatment experiments

Cohesin-complex droplets were formed on doubly tethered DNA molecules after 5 min injection of 10 nM Alexa647-cohesin complex. Then, a different salt buffer (50 mM Tris-pH7.5, 50/100/500 mM NaCl, 2.5 mM MgCl_2_, 250 nM SxO, and the gloxy oxygen scavenging buffer) was injected (Fig. 2, D and E). For the hexanediol experiment, 10 % or 20 % 1,6 hexanediol (Sigma-Aldrich) in a reaction buffer (50 mM Tris-pH7.5, 50 mM NaCl, 2.5 mM MgCl_2_, 250 nM SxO, and the oxygen scavenging system) was injected. The decrease of the Alexa647-cohesin droplet intensities was monitored using a 642 nm laser (Fig. 2, E and G).

#### Image analysis and quantification

Immobilized λDNA fluorescence intensity profiles showing DNA clusters were obtained from the summation of the intensity values of 11 pixels obtained from a line perpendicular to the extended DNA in each frame. Background intensity was subtracted using a 2D-median smoothing. Afterwards, to correct intensity fluctuations such as bleaching, the intensities were normalized by the maximum value during the measurements. The kymograph was constructed after obtaining the intensities for all frames the normalized intensity profiles.

For the kinetics analysis of DNA cluster formation, we defined the region of interest (ROI) of both the clustered area and the area of the entire DNA using Image J (Fig. 1C left). Using ImageJ, we measured the sum of the intensities of the pixels in the ROI of each frame. Background intensities were subtracted. Using the intensity information, the size of DNA (in kbp) in the clustered area was obtained by normalization with the sum of the entire DNA intensity and multiplication by 48.5 kbp (*5*). To obtain the compaction time, we smoothed the time trace using the Savizky-Golay method with a moving window of 250 points. Then, we determine the compaction time between starting compaction point (5 %) and the ending point (95 %). To measure the kinetics of the cohesin-complex cluster formation and release, we used the ROI where the Alexa647-cohesin complex cluster was co-localized with the cluster formation, yielding intensity-time traces as shown in Fig. 1G. To measure the bleaching steps of a single Alexa647 cohesin complex (Fig. 1H inset), we did not use oxygen scavenging system (gloxy) to effectively bleach the Alexa647 fluorophores. To clearly obtain the step-size, we analyzed clusters whose initial intensity was comparably low, where only a limited number of steps (< 8) were observed. We applied a step-finding algorithm following a previously described algorithm (*32*).

Cohesin droplet diameter and circularity were measured by Fiji and Matlab (9) (Fig. 2, A-C). The cross-sectional intensity profiles of droplets were obtained by Fiji. The diameter was measured by the distance between two points at 20 % of the maximum fluorescence intensity of the Gaussian-fitted graph of cross-sectional intensity profiles of a droplet and using a home-built Matlab code (Fig. 2A). As a control, to show that our observed cohesin-complex/DNA droplets are larger than diffraction limit, 20 nm quantum dots (Qdot^TM^ 705, thermofisher) were non-specifically adsorbed on the slide glass and similarly analyzed. To extract the diameter of a single-quantum dot rather than QD clusters, we analyzed quantum dot fluorescence spots that showed blinking events. In order to show that the droplet is spherical, the circularity was measured using Fiji by applying a threshold of 20 % intensity of the average maximum intensity of the Gaussian fits of the droplets (Fig. 2C) (*9*).

#### Chromatin immunoprecipitation (ChIP) experiments using 1,6-hexanediol treatment of yeast cells

Cells were grown in Yeast-Peptone-Dextrose (YPD) medium to mid exponential phase before addition of 8 μg/mL nocodazole for two hours to achieve a mitotic arrest. A first aliquot of the culture was retrieved for ChIP analysis. 1,6-hexanediol was added at a concentration of 5%, together with 10 μg/ml digitonin to permeabilize the cell membrane for 10 minutes, conditions that did not impede cell growth or survival following washout. After taking a second aliquot, the culture was filtered, washed and resuspended in fresh medium containing nocodazole but no 1,6-hexanediol for recovery and the final aliquot was harvested after one hour. Chromatin immunoprecipitation followed a previously published protocol (*27*). Briefly, cells were fixed with formaldehyde and harvested. Protein extracts were prepared and chromatin disrupted by sonication. DNA fragments cross-linked to the protein of interest were enriched by immunoprecipitation. After reversal of cross-links, DNA both from immunoprecipitates and from whole cell extract was purified and quantified using the PowerUP SYBR Green Master Mix (ThermoFisher) and a Viia7 Real-Time PCR System (Thermo Fisher). The primer sequences used are listed in Table S1.

#### AFM imaging and image processing

To image cohesin-complex/DNA clusters, we mixed 10 nM cohesin complex and 4 ng/μL DNA of various length [0.1, 0.3, 0.5, 1, 3, 5, 10 kbp (Thermo Scientific), and 48.5 kbp (Promega)] in a reaction buffer (50 mM Tris-pH7.5, 50 mM NaCl, 2.5 mM MgCl_2_, 2mM TCEP) in an E-tube and incubated the mixture for 5 min (Fig. 1, J to M and Fig. 3, A to H). In the case of Figure 3H, to show single-cohesin complex mediated DNA bridging, we mixed 1 nM cohesin complex and 4 ng/μL 3kbp DNA and incubated them for a very short time (10 s). Afterwards, the mixtures were deposited onto mica that was pretreated with polyLysine (0.00001 %) (*33*). After briefly washing the sample using 3 mL milliQ water, the sample was dried using a nitrogen gun.

AFM measurements of the dried sample were performed on a Multimode AFM (Bruker), with a Nanoscope V controller and Nanoscope version 9.2 software. ScanAsyst-Air-HR cantilevers (Bruker, nominal stiffness and tip radius 0.4 N/m,and 2 nm, respectively) were used. PeakForce Tapping mode was used with an 8 kHz oscillation frequency, and a peak force setpoint value less than 70 pN was used to minimize sample invasiveness caused by sample and tip interaction. For imaging the proteins and protein/DNA mixtures, the scan area of 10 μm × 10 μm with 5,120 × 5,120 pixels were recorded at the scanning speed of 0.7 Hz. All measurements were performed at room temperature.

For image processing of dry AFM images, Gwyddion version 2.53 was used. First, background was subtracted, and transient noise was filtered. To ensure that only the empty surface was used for background subtraction, masking particles and subtracting (planar and/or line-by-line) background polynomials was employed by excluding the masked regions. Horizontal scars, which occasionally occur because of feedback instabilities or protein sticking to the AFM tip, were removed. Afterwards, plane background subtraction was applied. Finally, the blind tip estimation was used to estimate the shape of the tip, and surface reconstruction was done to reduce the broadened effects caused by tip convolution (the widening of images induced because the AFM tip size is not zero)(*34*).

To measure the volume of a single protein (fig. S3), using Gwyddion, we masked each protein on the images and we performed grain volume measurement to obtain the volume information of the each masked protein For the cluster volume measurements, the territory of the proteins areas bound to DNA was defined by Gwyddion manually. The volume of the defined area was obtained by Gwyddion. To analyze the volume of the cohesin-complex/DNA clusters, to obtain higher than 99.9 % confidence level, seven standard deviation plus the average of the volume of a single cohesin-complex was used as a threshold (2,134 nm^3^). The defined area whose cluster volume is higher than the threshold was considered as a cluster.

### Supplementary Text

#### Theoretical calculation of a critical looping length

The elastic bending energy of a DNA segment of length *l* is

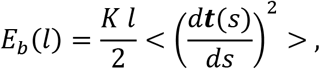

where ***t***(*s*) is the tangent to the DNA at point *s, K* is the elastic modulus which equals *K = k_B_T l_p_*, with *l_p_* the persistence length of DNA in nm, i.e., *l_p_* = 50 nm, and <> denotes averaging over the DNA segments *l*. For a perfect circle of radius R, the curvature is d***t***(*s*)/*ds* = 1/*R* and hence

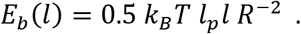

Using this equation, we may ask what is the critical length at which thermal fluctuations are able to deform the segment so that it can loop onto itself, approximately the shape of a circle. For this, we must solve *E_b_*(*l*\*) = *k_B_T* or

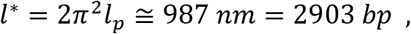

where we used 2*πR = I*\*. This means that a segment 3 kbp long can loop onto itself due to thermal fluctuations, whereas a loop on a fragment, e.g., 100 bp, requires an energy of about 30 *k_B_T* and is therefore very rare to observe.

Another way to compute the typical size of loops observed in freely fluctuating DNA is by balancing the energy of bending with the (loss of) entropy due to looping. One should then minimize the following free energy

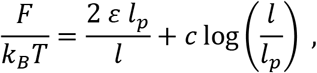

where the first term is now the bending energy of a generalized shape that is not strictly a perfect circle but can have a kink. This freedom is parametrized by the parameter *ε*. The second term is the entropic loss associated with a loop of size *l*, computed as —*k_B_T*log[(*l/l_p_*)^-*c*^]. The exponent *c* characterizes the contact probability of two segments in a polymer (e.g., *c* = 1.5 for an ideal random walk). Setting *ε* = 16, valid for a tear-drop shape (*35*), we obtain that the minimum of the free energy is at *l*\* = 3.2 kbp.

Both calculations suggest that one requires DNA segments longer than about 3 kbp in order to observe looping and hence bridging-induced clustering/phase separation whereby a stochastic thermal loop stabilized by a cohesin-complex can act as a nucleation point for clustering. The clustering is entropically and energetically favored over a dispersed situation because once a loop is formed, many cohesin complexes can bridge DNA without the need of forming new loops (that cost energy and entropy). This positive feedback – looping attracts bridge proteins that drive/stabilize further looping – is known as bridging induced attraction (*17, 18*).

#### Modelling of DNA and cohesin

We modelled DNA as a semiflexible polymer made of beads of size *σ* = 2.5 nm = 7.3 bp and with persistence length *l_p_* = 20*σ* = 50 nm = 150 bp. The protein bridges are made by a spherical central bead of size *σ_b_* = 25 nm which is decorated by 2, or 3 patches or made fully attractive (‘multi’). The patches (or the body itself in absence of patches) act as DNA binding site for the protein as described below.

In more detail, the excluded volume interactions between DNA beads (including consecutive ones along the DNA), obeyed the shifted and truncated Lennard-Jones (LJ) potential

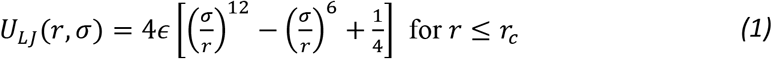

and 0 otherwise. In this equation, *r* denotes the separation between the bead centers. The cutoff distance *r_c_* = 2^1/6^ *σ* was chosen so that only the repulsive part of the Lennard-Jones is used for DNA beads, i.e., there is no attraction. Consecutive monomers along the DNA contour length were connected by the finitely extensible nonlinear elastic (FENE) potential, given by

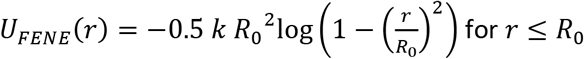

and infinity otherwise. In this equation *k* = 30 *ε/σ*^2^ is the spring constant and *R*_0_ = 1.5 *σ* is the maximum extension of the FENE bond. The persistence length for the DNA was introduced by adding a bending energy penalty between triplets of consecutive beads along the DNA as

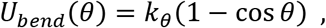

where *θ* is the angle formed between adjacent bonds, i.e. cos^-1^(***t***_*l*_ ***t***_*l*+1_/|***t**_i_* ||***t***_*l*+1_|) with ***t***_*l*_ the tangent at segment *i*, and *k_θ_* = 20 *k_B_T* the bending constant.

The excluded volume interactions between cohesin proteins were accounted for through the steric repulsion between the central spherical beads forming the body of the bridge proteins. In this case, the shifted and truncated Lennard-Jones (LJ) potential (Eq. (1)) was employed again as *U_LJ_*(*r, σ_b_*), where *σ_b_* = 25 nm and this time the cutoff distance was *r_c_* = 2^1/6^ *σb*, chosen so that only the repulsive part of the LJ potential was used for the central part of the cohesin-complex.

The interaction between DNA and the central body of the model cohesin was also captured by a repulsive LJ potential where the unit length is now (*σ* + *σ_b_*)/2 and the cut-off was chosen as r_c_ = 2^1/6^ (*σ* + *σ_b_*)/2 for bi- and tri-point interactions while as r_c_ = 1.8 (*σ* + *σ_b_*)/2 for multi-point interactions. The former retains only the repulsive part of the potential while the latter choice includes a spherical shell in which the spherical body is attractive to DNA beads with a binding energy that we set to *ε* = 0.7 *k_B_T*. These choices are well-established in the field for modelling DNA and chromatin binding proteins such as transcription factors (*18, 36*).

To model bi-and tri-point bridging we decorated the central spherical bead with two (for bi-point) and three (for tri-point) patches (see fig. S7, A to C). These patches interacted with DNA beads through the following Morse potential

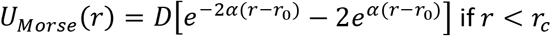

and 0 otherwise. The parameters of the potential were *D* = 40 *k_B_T, α* = 5*σ*^-1^, *r*_0_ = 0 and *r_c_* = 3 *σ*. These parameters ensured that the attraction is strong, but very short ranged. The choice of the equilibrium distance *r*_0_ = 0 was such that when a patch is bound to the DNA, it did sit at the center of the bead and excludes the binding of other patches as different patches interact via a repulsive-only LJ potential with size *σ.*

#### Molecular Dynamics Simulations

The static and kinetic properties of the systems are studied using fixed-volume and constant-temperature (NVT) Molecular Dynamics (MD) simulations with implicit solvent and periodic boundary conditions. MD simulations were performed using the LAMMPS package [http://lammps.sandia.gov/]. The equations of motion were integrated using a velocity-Verlet algorithm, in which all beads are weakly coupled to a Langevin heat bath with a local damping constant set to *γ =* 1 so that the inertial time equals the Lennard-Jones time 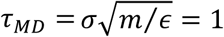 and *m* = 1 is the mass of the beads. The integration time step was set to δ*t* = 0.01 *τ_MD_*.

The bi- and tri-point bridging cohesin-complexes (modelled as a spherical body and patches) are evolved as rigid bodies with *γ* = 3 and *γ* = 4, respectively. For short DNAs (less than 10 kbp), we initialized the polymers as random walks and let them equilibrate with only repulsive conditions for long enough to swell the DNA, before turning on the attraction between proteins and DNA and proceed with the compaction. For longer DNAs, starting from a random walk yielded too heavily knotted/entangled conformations which were too slow to relax. Therefore we started from a helicoidal conformation and let equilibrate for 10^5^*τ_MD_*. For 40 kbp DNA, we equilibrated for 10^6^*τ_MD_*.

We monitored the radius of gyration defined as

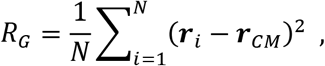

where *N* is the number of beads in the polymer and ***r**_CM_* is the position of its centre of mass. The final averages are performed over the last 10% of the time series (typically 500 frames spanning 5 10^4^*τ_MD_* time steps) in steady state and across at least 10 independent replicas.

Snapshots from equilibrated states are shown in fig. S7, A to C and Movie S5, whereas the radius of gyration as a function of DNA length is shown in fig. S7, D and E. As one can notice, the best agreement with the AFM scaling data is given by the case in which cohesin is modelled as a bi-point bridging protein – consistent with AFM images showing cohesin that was bridging DNA in 2 points. While DNA bound by many point bridges yielded a collapsed globule (*R_G_* ~ *L^α^* with *α* = 1/3), binding proteins with fewer binding sites are common *in vivo* but under-explored in the computational literature. Here we show that these may be leading to a qualitatively different DNA and chromatin folding with respect to their multipoint bridging counterparts.

#### *In silico* HiC Maps

To study how bi- and tri-point bridging proteins may affect the large-scale folding of DNA/chromatin *in vivo,* we performed simulations of DNA modelled as described above, i.e., *σ* = 2.5 nm, *l_p_* = 20 *σ, L* = 1000 beads in dilute conditions and with 100 cohesin proteins of size *σ_b_* = 2.5 nm, which could be either multi-, bi-, or tri-point bridging. DNA was patterned with alternating sticky or non-sticky blocks of length 100 beads. This “blockcopolymer-like” set up is intended to highlight the emergence of compartments in the contact map which is obtained *in silico* by calculating the frequency with which any two beads (*i, j*) are ‘in contact’, i.e., within a distance *d = 3σ* = 7.5 nm. A contact map was created by averaging the entries of the contact matrix over a short time window in steady state and across at least 20 independent replicas. The maps were then normalized so that the sum of all entries equals 1. The results compared for bi-, tri- and multi-bridging are shown in fig. S8, A and B.

**Table S1.**
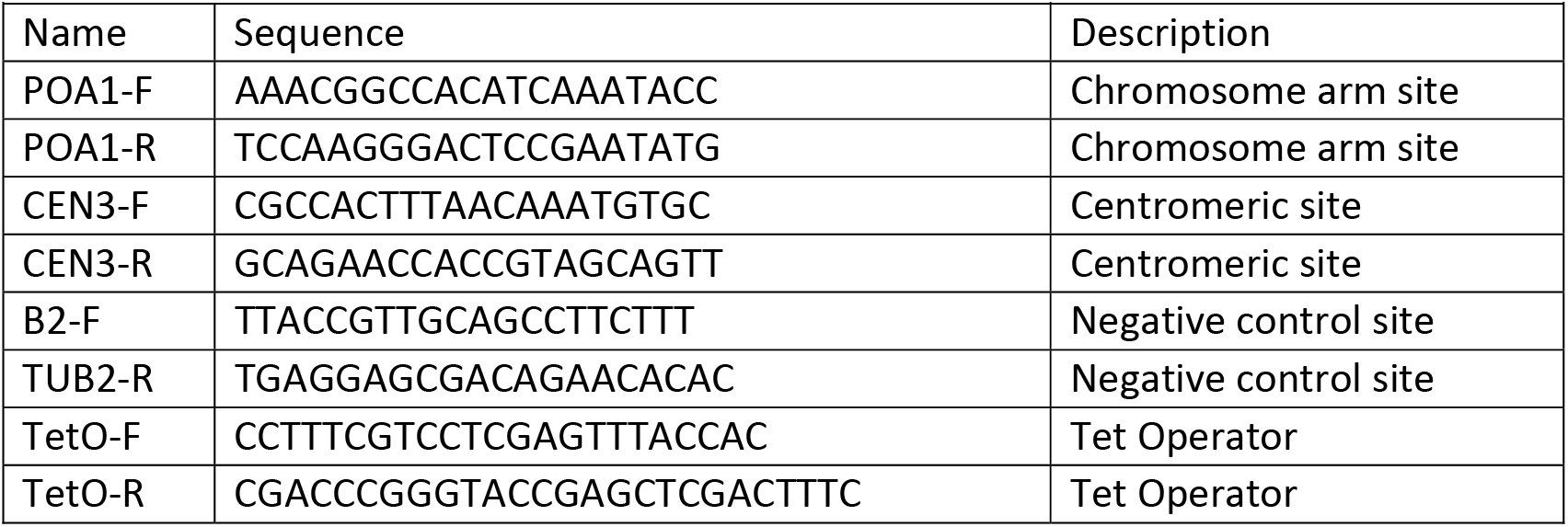
Primer pairs used for Chip-qPCR analyses

**Fig. S1.**
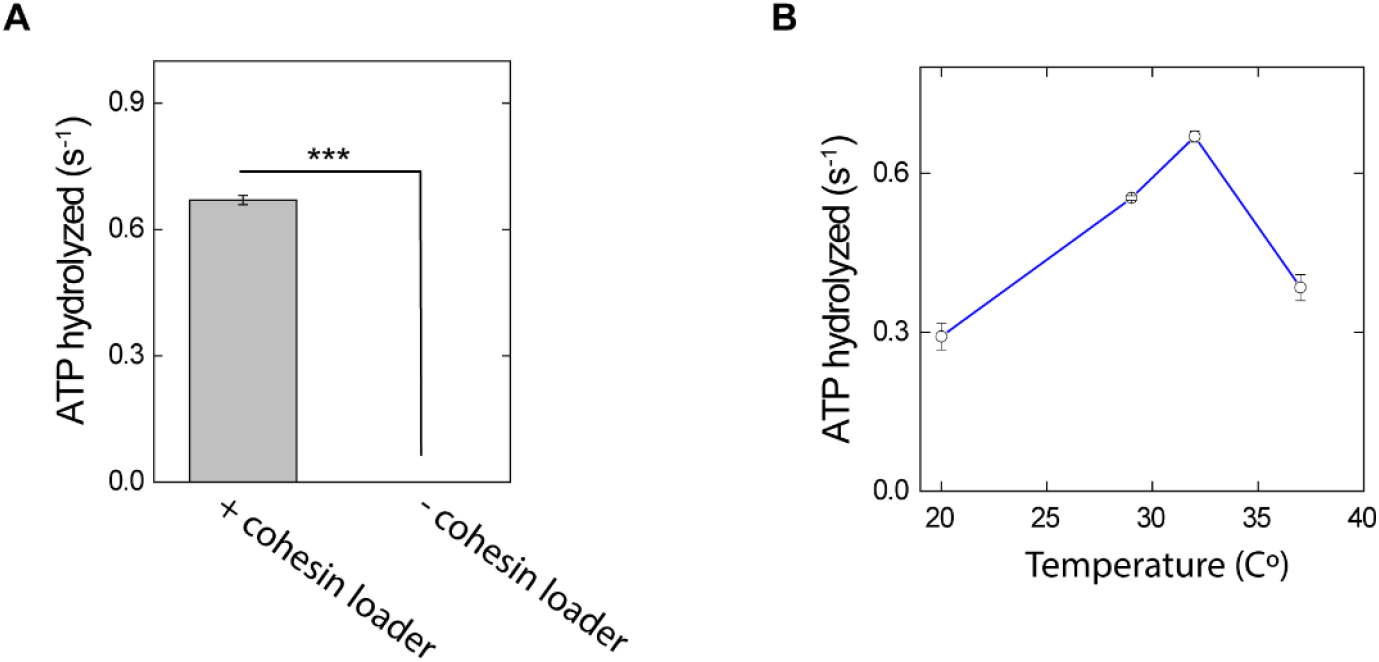
ATPase activity test of cohesin. **(A)** ATP hydrolysis rate with/without cohesin loader at 32 C° (*N* = 3). **(B)** ATPase activity versus temperature (*N* = 3). Errors are SD. *** indicates *P* < 0.001, assessed using the two-tailed Student’s *t*-test.

**Fig. S2.**
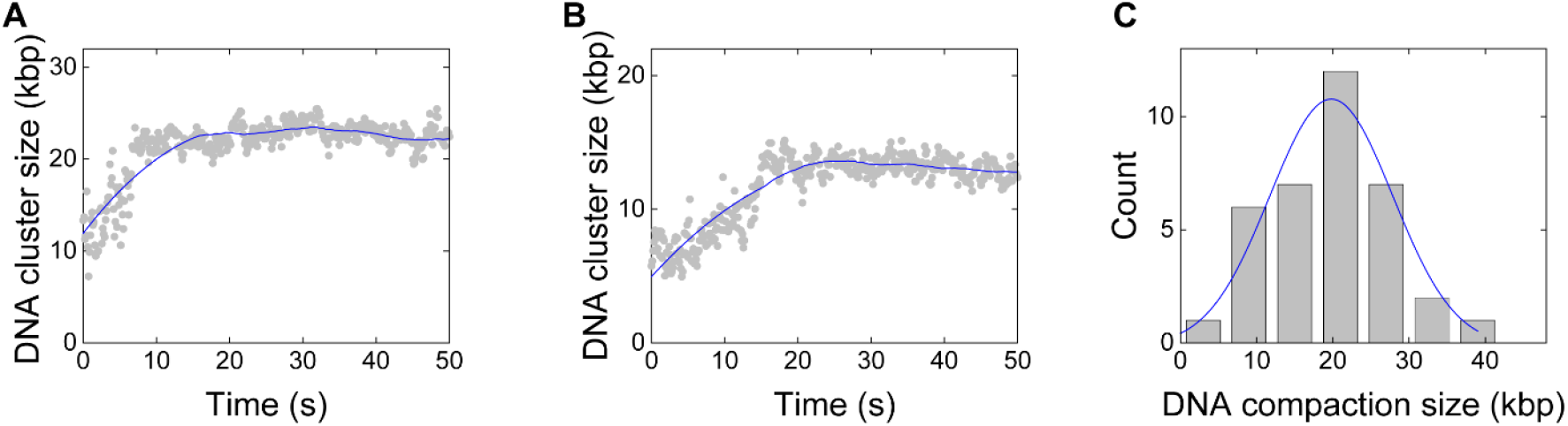
DNA cluster size kinetics. (A and B) Representative traces of DNA cluster formation. **(C)** DNA compaction size at saturation. Blue line is a Gaussian fit yielding 20 ± 8 kbp (mean ± SD) (*N* = 36)

**Fig. S3.**
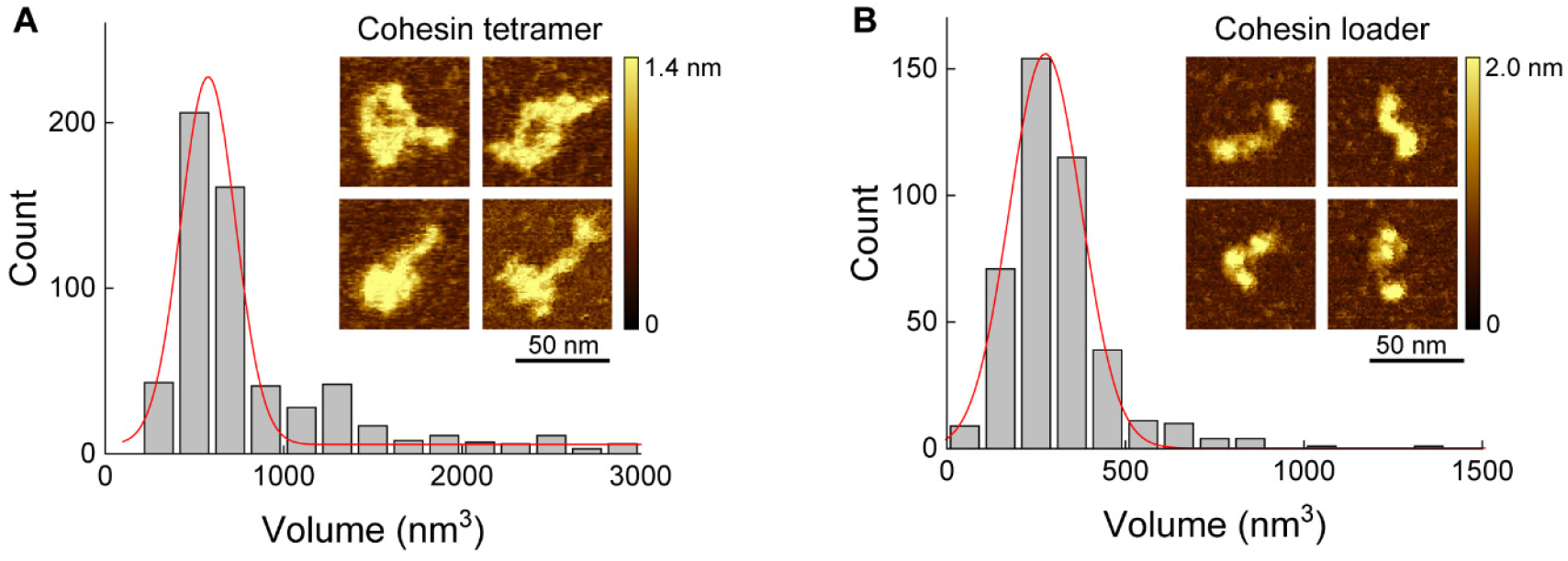
Volume of individual cohesin teramers (A) and cohesin loaders (B). Red lines indicate Gaussian fits (mean ± SD = 577 ± 153 nm^3^ and 276 ± 101 nm^3^, *N* = 602 and 419, for cohesin tetramer and cohesin loader, respectively). Representative AFM images are shown in insets.

**Fig. S4.**
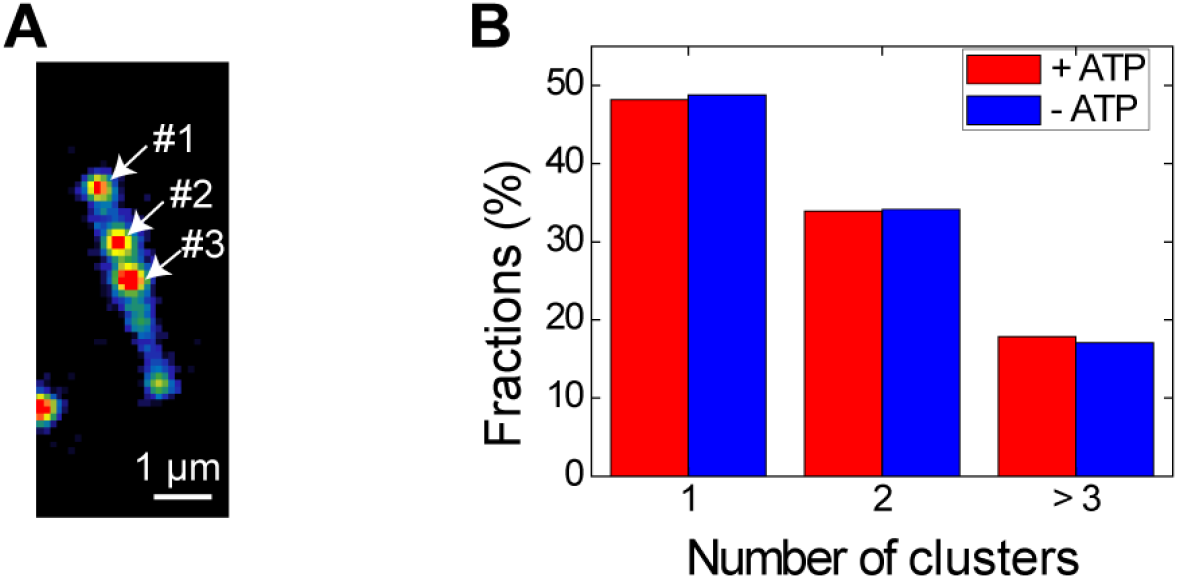
Number of DNA clusters on a DNA molecule after cohesin-complex injection. (A) Representative snapshot of a DNA molecule with three clusters. (B) Fractions of DNA with different number of clusters (*N* = 64 and 53 for experiments with/without ATP).

**Fig. S5.**
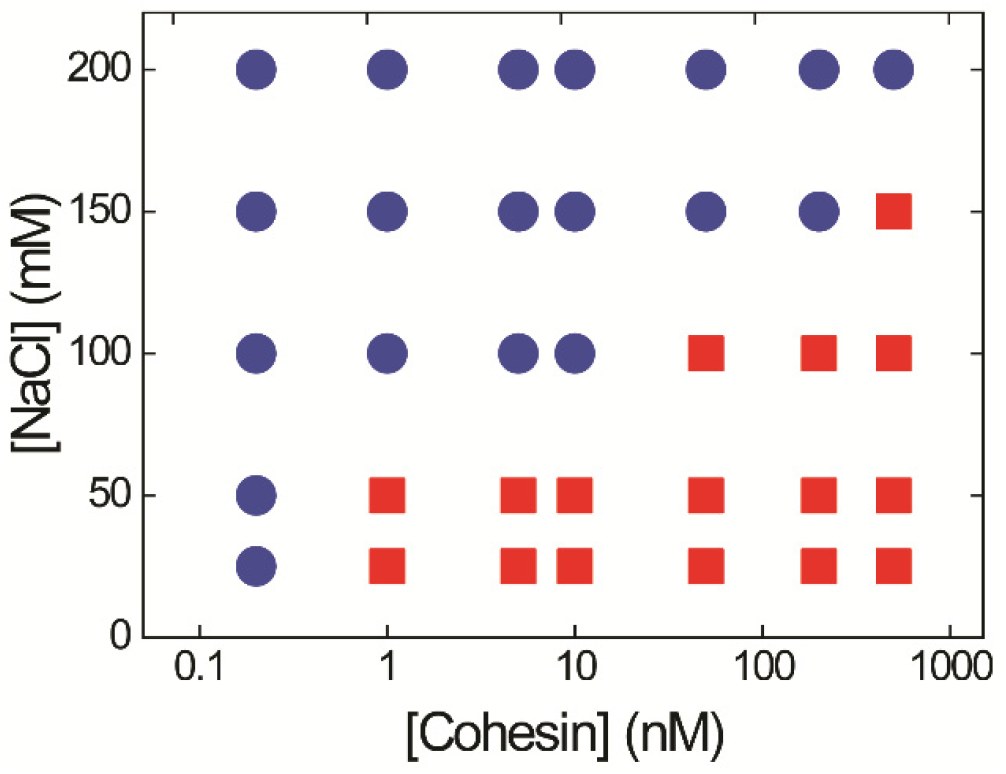
Phase-diagram of cluster formation induced by cohesin complex and λDNA at different salt and cohesin-complex concentrations, obtained at higher concentration of SxO (250 nM). A similar phase diagram is obtained as compared to Fig. 2F, albeit with a slight shift in the boundary line between phase separation and no phase separation. A higher SxO concentration may slightly inhibit the protein-DNA interactions (*37*).

**Fig. S6.**
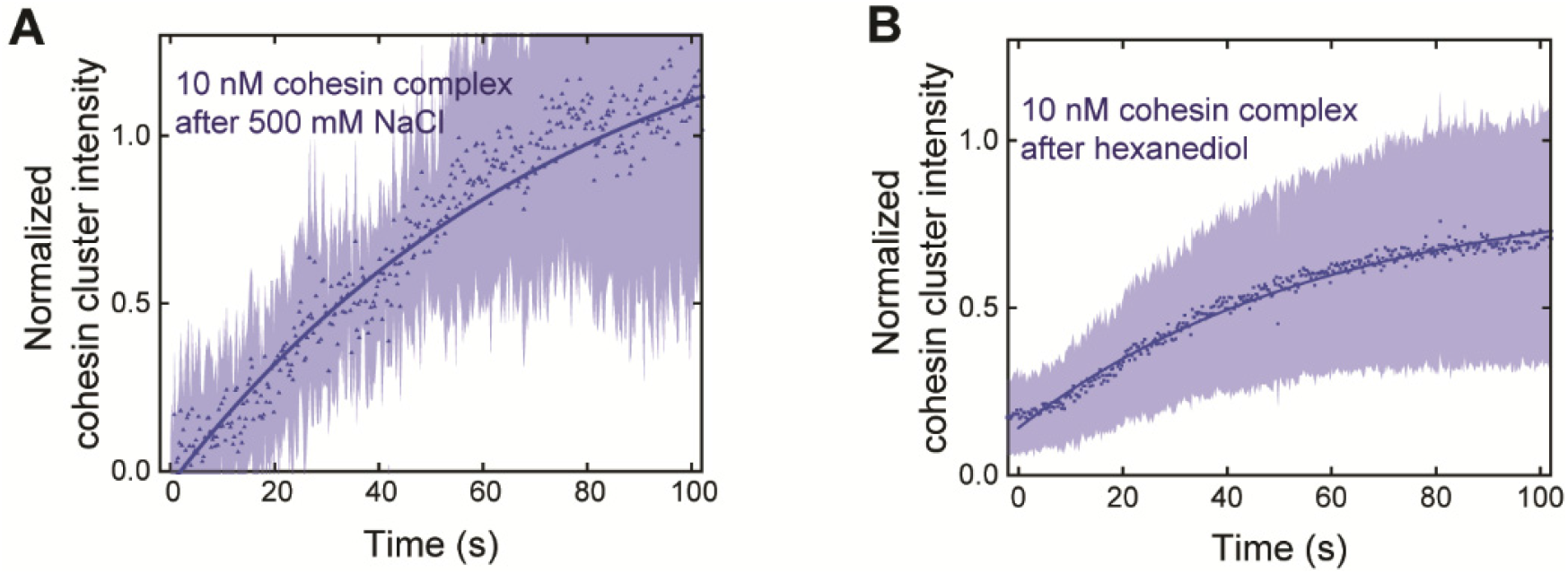
Recovery experiments. **(A)** After 500 mM NaCl washing of the cohesin-complex/DNA clusters (i.e. Fig. 2H), 10 nM labeled cohesin complex was reintroduced, and clusters reappeared. **(B)** After 20 % 1,6-hexanediol washing of cohesin-complex/DNA cluster (i.e. Fig. 2J), 10 nM labeled cohesin complex was reintroduced and clusters reappeared. Solid lines t show exponential fits, 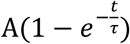 and light-colored areas denote the SD of the data points. (recovery time: 43.6 s and 58.3 s after 500 mM NaCl (A) and 20 % 1,6-hexanediol buffer wash (B), respectively; *N* = 60 and 40).

**Fig. S7.**
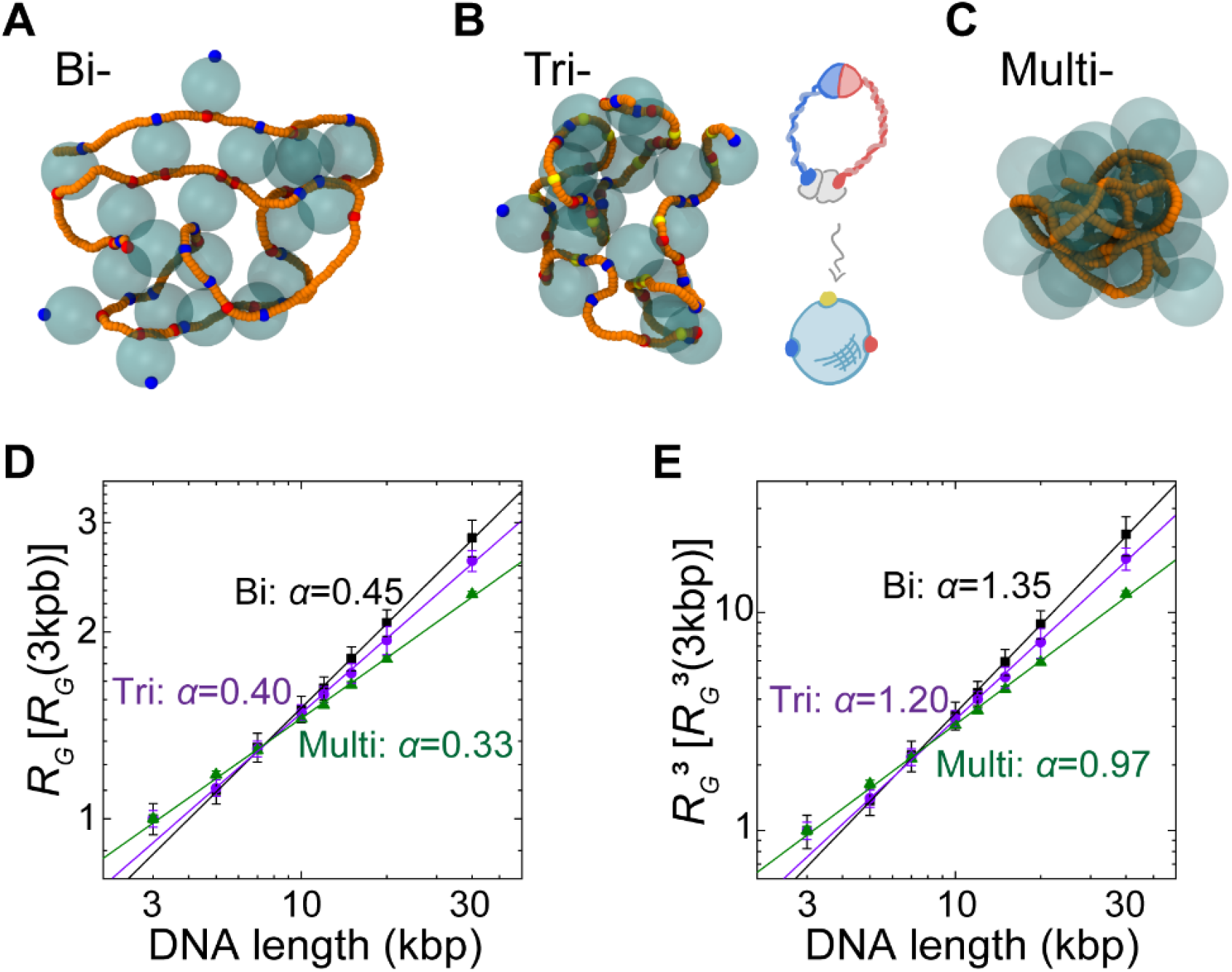
MD simulations of cohesin-mediated DNA cluster formation via different numbers of connectivity. **(A-C)** Snapshots from MD simulations of equilibrated conformations of DNA (orange) and cohesin-complexes modelled as spheres (cyan, transparent) with (A) two (bi-bridging, blue, red, Movie S5) (B) three (tri-bridging, yellow, blue, red) and (C) multi-bridging. A sketch showing the modelling of cohesin as a tri-point bridge protein is also shown. (**D**) Scaling of the radius of gyration *R_G_*, normalized by its value at 3 kbp as a function of total DNA length. Error bars are s.e.m. for each point. We observe *R_G_* ≅ *l^α^* with *α* ≅ 0.45 ± 0.01 for bibridges (black, similar to the scaling of AFM data, see main text), while *α* ≅ 0.40 ± 0.01 for tri-point bridges (violet), and *α* ≅ 0.33 ± 0.01 for the multi-point bridges (green). Errors are SD. **(E)** Scaling of 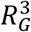 (cluster volume) normalized by its value at 3 kbp. Error bars are SEM. for each point. These plots suggest that the bi-point bridging is the one that best recapitulates the scaling observed in the AFM data, i.e., *α* ≅ 1.35 ± 0.02 (bi), whereas *α* ≅ 1.20 ± 0.02(tri) and *α* ≅ 0.97 ± 0.02 (multi). Errors are SD.

**Fig. S8.**
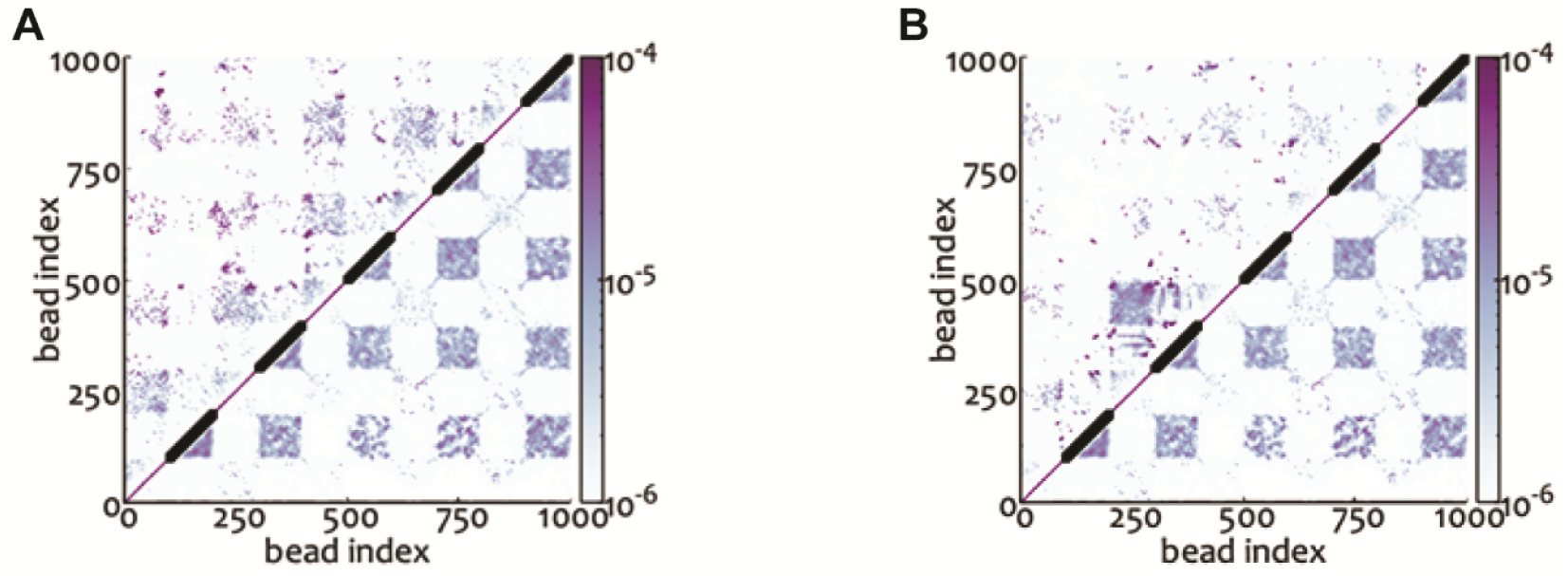
*In silico* Hi-C maps from simulations, obtained for different connectivities. **(A)** Top triangle: bi-point bridges. Bottom triangle: multi-point bridges. **(B)** Top triangle: tri-point bridges. Bottom triangle: multi-point bridges. The intensity of the color map at site (i,j) is computed as the probability (averaged over time and ensemble) of segments i and j to be in contact (defined as being less than 3σ apart). Furthermore, each entry of each map is divided by a normalization factor so that the sum of all the entries of the matrix sums to 1. Black thick segments along the diagonal show the stretches of DNA that are made sticky for the proteins.

**Fig. S9.**
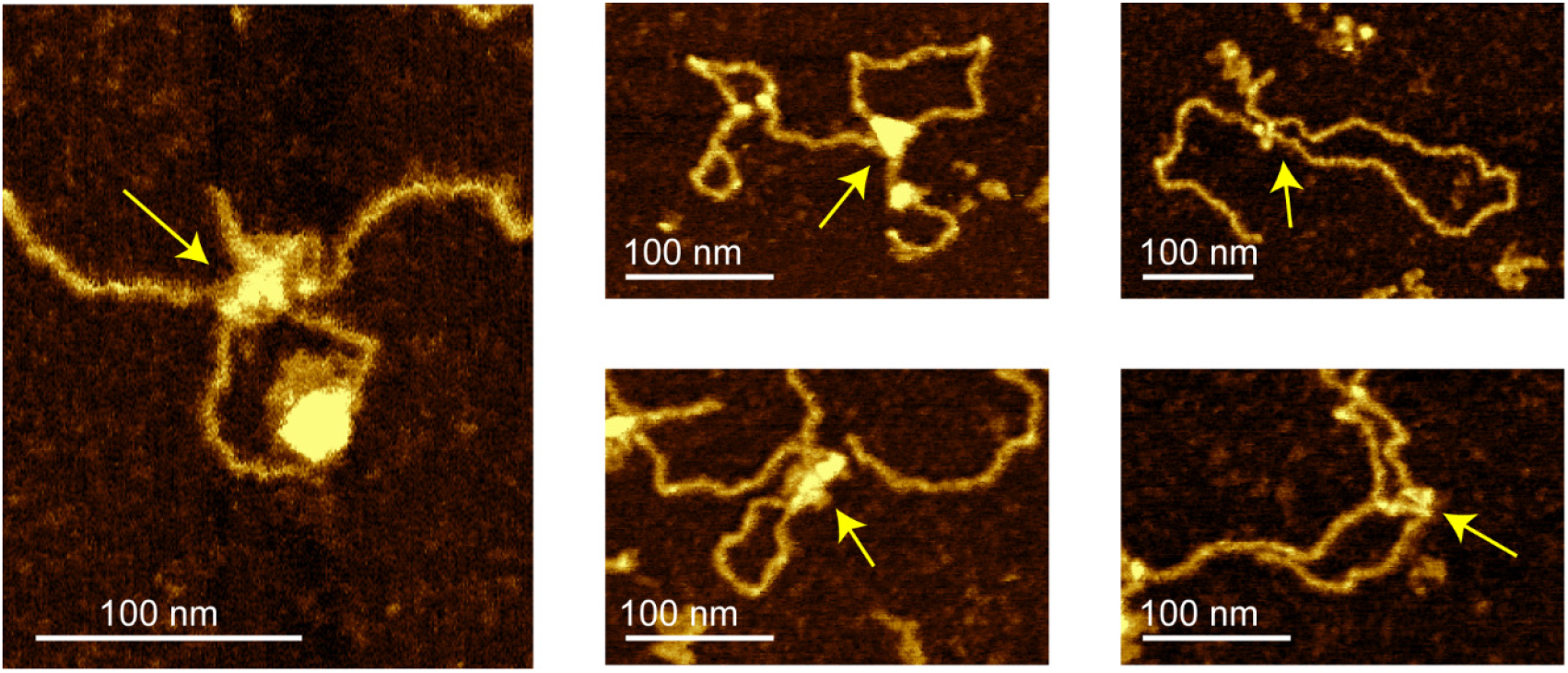
Further examples of AFM images where a single cohesin complex bridges DNA. Yellow arrows indicate bridged DNA regions.

### Supplementary movies legends

**Movie S1.** Movie showing cohesin-complex induced DNA cluster formation along a double tethered SxO-stained DNA.

**Movie S2.** Movie showing a side-flow experiment with a constant flow where cohesin-complex is injected. Initial frames shows an inverted U shape because of the side flow. A bright fluorescence spot grows to form a large cluster.

**Movie S3.** Movie showing both SxO-stained DNA (green) and Alexa647-labeled cohesin-complex (red), showing the correlative cluster spots of Alexa647-cohesin complex and DNA.

**Movie S4.** Movie of overlaid SxO-stained channel (green) and Alexa647-labeled cohesin complex (red), showing that two cohesin clusters do merge over time.

**Movie S5**. MD simulation of cohesin-driven clustering with DNA. Left: Folding process for a 20kb-long DNA driven by bi-bridging proteins. The DNA is shown in orange while proteins are shown, to scale, as transparent spheres (cyan) with blue-red patches that stick to the DNA. Notice that in steady state the DNA is not fully crumpled but remains partly open. Right: Same for multi-bridging proteins. Here, in steady state, the DNA becomes a globule, as one would expect for a collapsed polymer. Initial conditions were chosen from an equilibrated simulation in purely self-avoiding conditions.

